# Memory and Perception-based Facial Image Reconstruction

**DOI:** 10.1101/122705

**Authors:** Chi-Hsun Chang, Dan Nemrodov, Andy C. H. Lee, Adrian Nestor

**Affiliations:** Department of Psychology at Scarborough, University of Toronto, Toronto, Ontario, Canada; Rotman Research Institute, Baycrest Centre, Toronto, Ontario, Canada

## Abstract

Visual memory for faces has been extensively researched, especially regarding the main factors that influence face memorability. However, what we remember exactly about a face, namely, the pictorial content of visual memory, remains largely unclear. The current work aims to elucidate this issue by reconstructing face images from both perceptual and memory-based behavioural data. Specifically, our work builds upon and further validates the hypothesis that visual memory and perception share a common representational basis underlying facial identity recognition. To this end, we derived facial features directly from perceptual data and then used such features for image reconstruction separately from perception and memory data. Successful levels of reconstruction were achieved in both cases for newly-learned faces as well as for familiar faces retrieved from long-term memory. Theoretically, this work provides insights into the content of memory-based representations while, practically, it opens the path to novel applications, such as computer-based ‘sketch artists’.

---

Remembering the visual appearance of a known face is a crucial part of everyday life. To date, extensive research has established the impact of specific contextual and intrinsic facial properties on face memorability (e.g., distinctiveness, familiarity, inter-group similarity, race, emotional expression, and trustworthiness, to name a few)^1–9^. Yet, much less is currently known about the concrete pictorial information associated with retrieving a face from memory. Arguably, elucidating this issue can provide valuable insights into the nature of the representations subserving face memory and also, into their relationship with face perception.

Accordingly, the current work seeks to elucidate the representational content of visual face memory through the novel use of image reconstruction. Previously, reconstruction approaches have been mainly directed at estimating the perceptual representations of an observer from patterns of neural activation^10–14^. Importantly though, reconstruction has not targeted longterm memory and its pictorial content as derived from behavioural data (but see recent work on neural-based image reconstruction from working memory^15^). To handle this challenge, here, we appeal to a robust reconstruction approach^12^ that capitalises on the structure of internal representations as reflected by empirical data irrespective of their modality (e.g., neural or behavioural). Further, this approach has a twofold goal of deriving facial features directly from empirical data and then using them in the process of image reconstruction.

Theoretically, at the core of our work lies the concept of face space^16^, a multidimensional construct comprising a population of faces with the property that the distance between any pair of faces reflects their psychological similarity^17–20^. Critical for our purposes, perceptual face space and its memory-based counterpart may be closely related^21^ allowing, in theory, the use of the former to inform the latter. Accordingly, here we rely on behavioral estimates of face similarity, whether between pairs of stimuli or between a stimulus and a face recalled from memory, to construct an integrated perception-memory face space. This construct allows the derivation of perceptual features, namely global pixel image intensities rather than local face parts (e.g., an eye), that exploits its organisation through an analogue of reverse correlation^22–24^. Such features are then combined to deliver image reconstructions for a novel set of faces projected in this space. Naturally, this approach allows perceptual and memory-based reconstructions alike depending on whether the target faces are perceived or remembered – image reconstruction from memory is applied here both to novel faces, learned over the course of the experiment, and to famous faces retrieved from long-term memory. Of particular note, successful memory-based reconstruction from an integrated perception-memory face space would provide strong evidence for shared representations underlying face perception and face memory.

Finally, since subjective personal experience is likely to shape substantially an individual’s memory for faces^25^, the present work seeks proof of principle that reconstruction can be performed individually, rather than at the group level, provided that sufficient data is collected to allow a robust approximation of face representations in single participants. To handle this challenge, data subserving reconstruction purposes were collected, across multiple experimental sessions, for each of three participants (Experiment 1); then, the accuracy of individual-based reconstructions was assessed objectively with respect to image pixel intensities (Experiment 1) as well as experimentally by a larger group of participants (Experiment 2). From a translational standpoint, the current strategy carries significance in that practical applications of such methodology are likely to target single individual data (e.g., independent estimation and visualisation of face memory in single eyewitnesses). At the same time, reconstructed faces should be recognisable by most individuals sufficiently familiar with the intended targets – such individuals would include, by necessity, but not be limited to the individuals who provided reconstruction data.

In sum, the current work aims to provide a theoretical framework for integrating the study of perceptual and memory representations as well as new methodology for estimating the pictorial content of visual memory in single individuals.

## Results

### Reconstruction approach

Facial images, including three newly-learned faces, three famous faces retrieved from long-term memory, as well as the 57 unfamiliar faces perceived by participants, were reconstructed separately for each of three participants in Experiment 1 (NC, CB and SA). This endeavour was pursued through a sequence of steps that capitalised on the structure of face space for the purpose of feature derivation and image reconstruction. In short, this sequence included: (i) constructing a multidimensional face space (Fig. 1c) from experimental estimates of pairwise face similarity (Fig. 1a, b) using multidimensional scaling (MDS); (ii) deriving classification images (CIM) for each dimension and assessing their significance regarding the inclusion of relevant visual information (Fig. 1d); (iii) projecting the target face into face space (i.e., approximating its coordinates in that space); and (iv) reconstructing the target by combining significant CIMs proportionally with the target’s coordinates in face space (see Methods and Fig. 1 for the further details).

**Figure 1.**
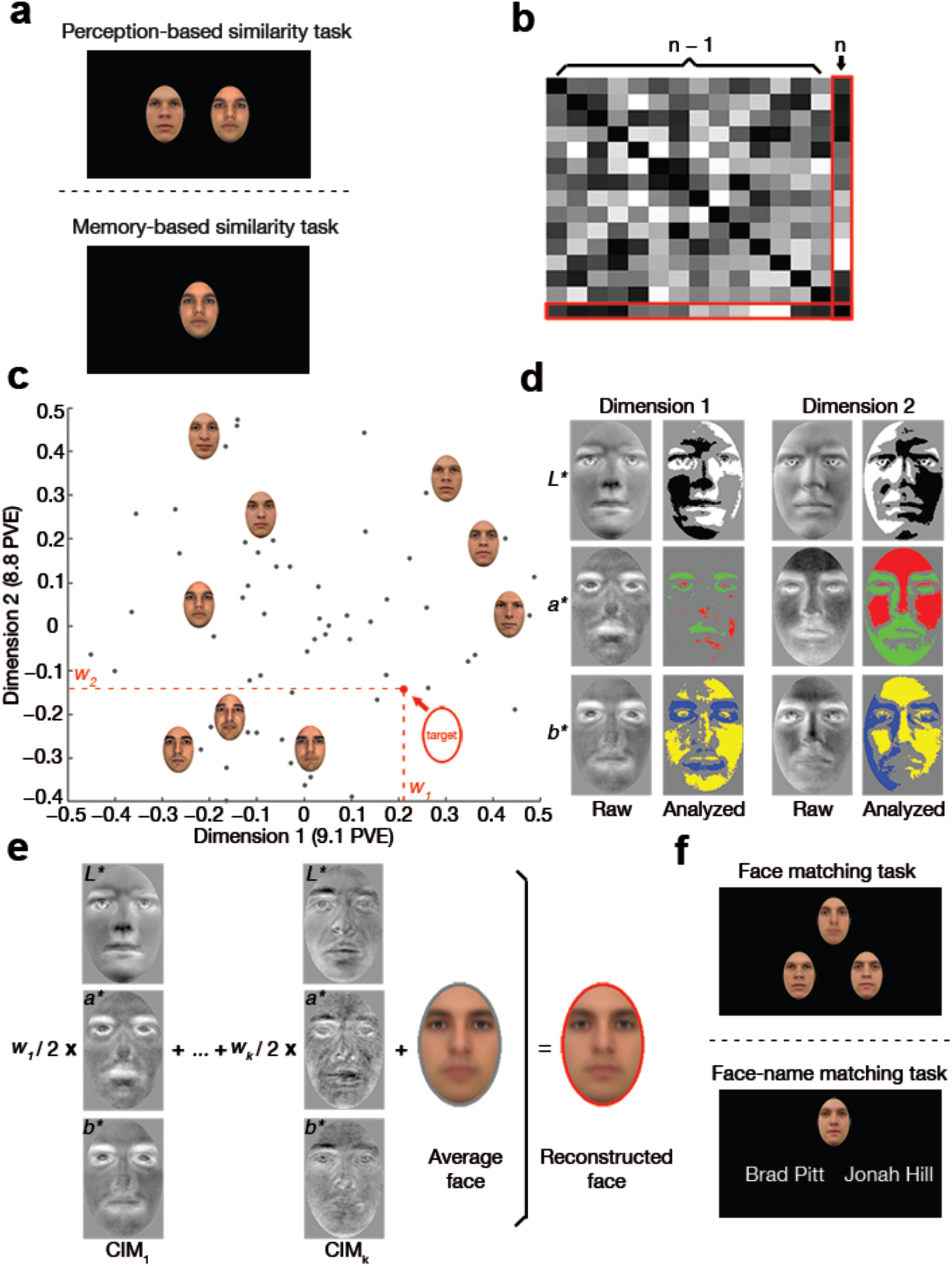
Reconstruction procedure and result evaluation: (a) participants rate the face similarity of two stimuli or of a single stimulus against a facial identity retrieved from memory; (b) pairwise face similarities are converted into a confusability matrix of ***n*** distinct facial identities, where face ***n*** is the target face for reconstruction purposes; (c) face space is estimated from the similarity of ***n-1*** different faces and the coordinates of the target face are approximated within that space (only 2 dimensions are displayed for convenience; PVE – percent variance explained); (d) visual features corresponding to each dimension are derived through image classification from ***n-1*** faces and analysed, separately for each colour channel in CIEL*a*b*, with a pixelwise permutation test (FDR-corrected across pixels; q<0.10); (e) visual features are linearly combined to estimate the visual appearance of the target face (CIM_k_ – classification images corresponding to dimension ***k***); (f) face reconstructions are evaluated in a two-alternative forced choice task with face pairs, for novel faces, or with name pairs, for famous faces. As an illustration, (c) and (d) show intermediary reconstruction results for participant NC. (Due to copyright restrictions all images of original face stimuli have been replaced with artificially generated face images.)

Representative examples of reconstructed images are shown in Fig. 2 for all three categories of faces (i.e., unfamiliar, learned, or famous). Overall, face reconstructions appear to capture the visual characteristics necessary for face identification in all conditions.

**Figure 2.**
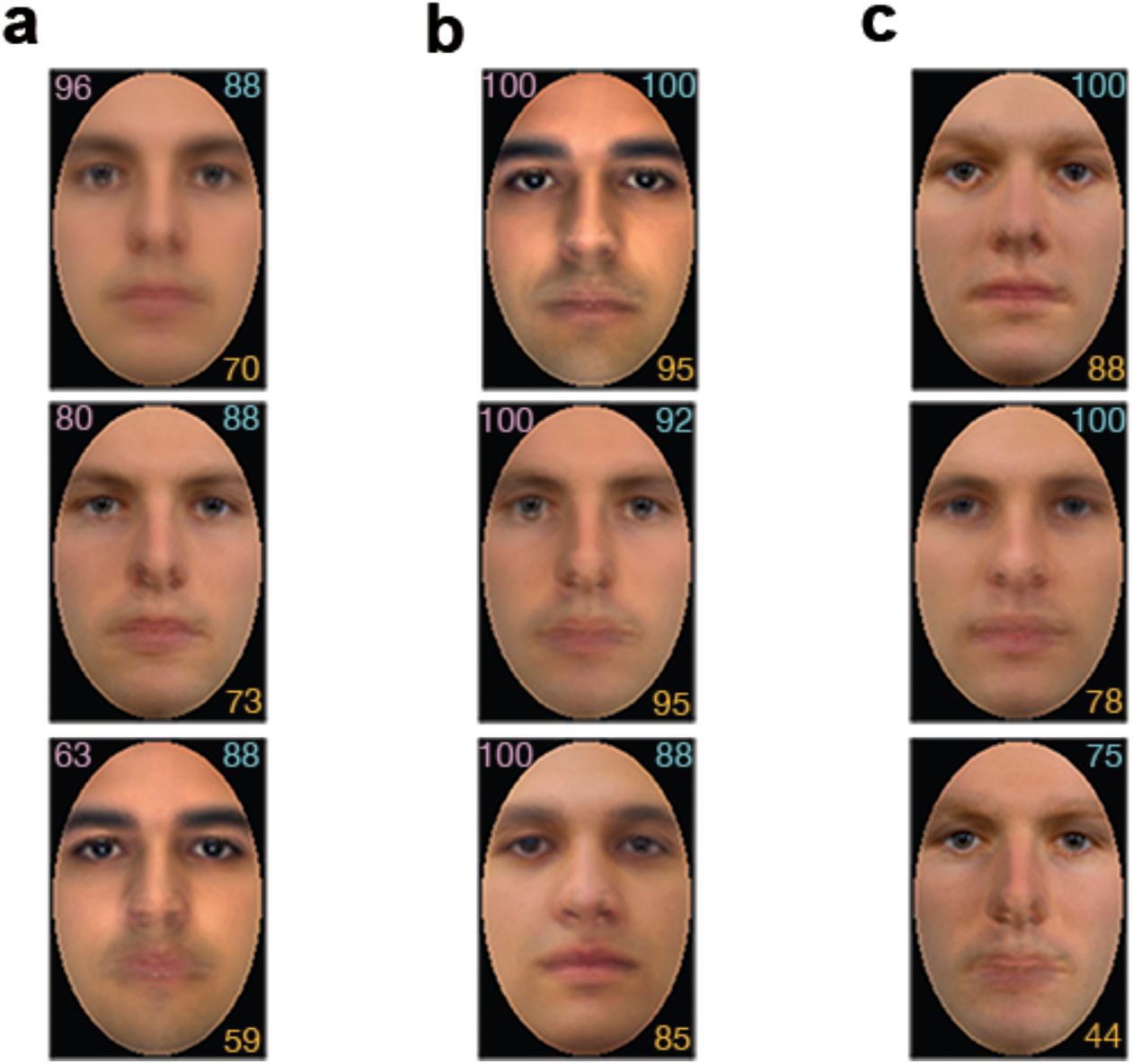
Examples of face reconstructions for participant NC. Results are shown separately for: (a) perceptual reconstructions, (b) memory-based reconstructions of learned faces and (c) memory-based reconstructions of famous faces (top to bottom: Jonah Hill, Chris Hemsworth, Jim Parsons) that participants were familiar with from their individual experience. Reconstruction accuracy (%) was estimated objectively through pixelwise image similarity (top left, a – b); additionally, accuracy was assessed experimentally by the same participant (top right) or by an independent group of naïve participants (bottom right) for all types of reconstruction. (Corresponding stimulus images for (a) and (b) could not be reproduced due to copyright restrictions.)

### Evaluation of reconstruction results

To assess reconstruction accuracy, an image-based evaluation procedure computed the pixelwise similarity between reconstructions and face images (e.g., actual stimuli). Then, the percentage of instances for which the reconstruction was more similar to its target than to any other image provided an estimate of image-based accuracy. An analogous, experimentally-based estimate, was further derived in Experiment 2 – a larger group of participants, including NC, CB and SA, were asked to judge the similarity between each reconstruction and two potential targets in a two-alternative forced-choice test.

Image-based estimates (Fig. 3a) as well as experiment-based estimates collected from the three participants above (Fig. 3b) as well as from other naïve participants (Fig. 3c) all confirmed that the reconstructions were successful. Of note, the average magnitude of reconstruction accuracy was above chance for every type of estimate, for every condition (i.e., perception, memory for learned faces and memory for famous faces), and for every set of reconstructions by participant (NC, CB and SA). Further statistical tests of perceptual reconstructions found that image-based and experimental-based accuracies computed for the three main participants were significant in all cases (comparisons against chance via two-tailed one-sample t-tests across stimuli,*ps* < .001) (see Table 1 and Fig. 3a, b).

**Figure 3.**
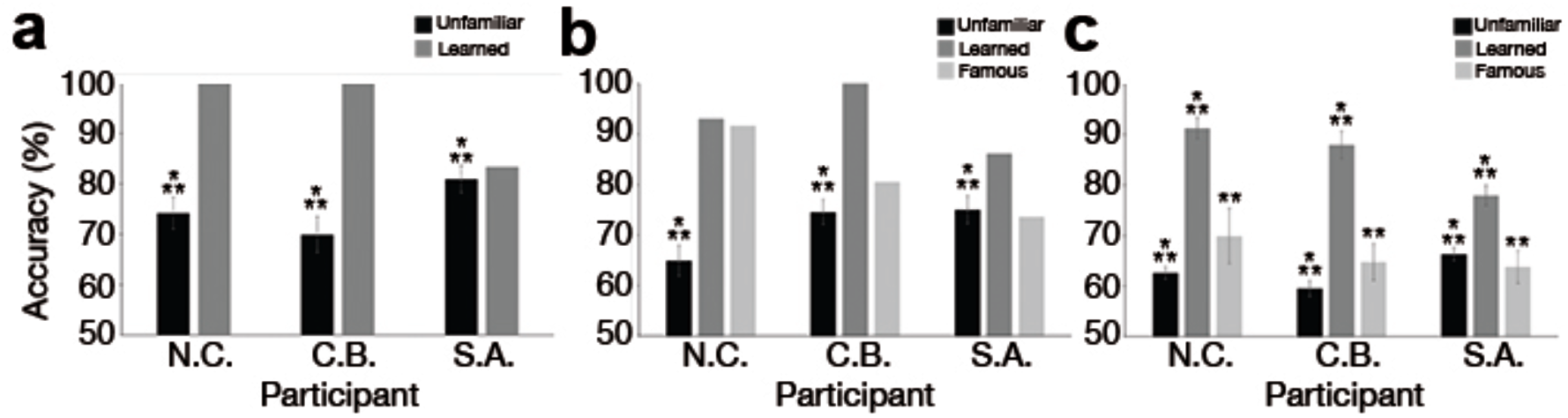
Average reconstruction accuracy for three participants (NC, CB and SA). Accuracy was estimated via (a) objective pixelwise similarity; (b) experimental data from the same participants, tested on their own reconstructions, and (c) experimental data from an independent group of participants (error bars show 1SE; ** - p<0.01 *** - p<0.001).

**Table 1.** Evaluation of perception-based reconstructions across stimuli

| Participants | Image-based accuracy |  |  | Experimentally-based accuracy |  |  |
| --- | --- | --- | --- | --- | --- | --- |
| | $t(56)$ | 95% CI | Cohen's $d$ | $t(56)$ | 95% CI | Cohen's $d$ |
| NC | 7.69 | [0.68, 0.80] | 1.02 | 4.88 | [0.59, 0.71] | 0.65 |
| CB | 5.60 | [0.63, 0.77] | 0.74 | 9.97 | [0.70, 0.80] | 1.32 |
| SA | 11.41 | [0.75, 0.86] | 1.51 | 8.93 | [0.69, 0.81] | 1.18 |

Note: CI = confidence interval.

To estimate more thoroughly reconstruction results, a two-way mixed-design analysis of variance (3 within-participants reconstruction types: unfamiliar, learned, or famous x 3 between-participants face triplets: NC, CB or SA) was applied to naïve participant data from Experiment 2. This analysis found a main effect of reconstruction type (F(1.289, 34.79) = 61.13, *p* < .001, η^2^ = .694, Greenhouse-Geiser correction for sphericity) and an interaction effect (F(2.577, 34.79) = 3.21, *p* = .04, η^2^ = .192, Greenhouse-Geiser correction), but no effect of face triplet. Further pairwise comparisons revealed that the accuracy of learned faces was significantly larger than that of either unfamiliar (t(29) = 12.44, *p* < .001, CI of the difference: [0.20, 0.26], *d* = 2.27) or famous faces (t(29) = 7.56,*p* < .001, CI: [0.13, 0.26], *d* = 1.38).

Importantly, comparisons against chance found that reconstructions were significant in all cases (two-tailed one-sample t-tests across participants; famous face reconstructions for NC, CB, and SA: *p* = .005, *p* = .003, *p* = .002, respectively; all other ps < .001) (see Fig. 3c and Table 2).

**Table 2.** Evaluation of reconstruction results across independent participants (Experiment 2)

| Participants | Reconstruction types | $t(9)$ | 95% CI | Cohen's $d$ |
| --- | --- | --- | --- | --- |
| NC | Perception/Unfamiliar | 10.22 | [0.60, 0.65] | 3.23 |
|  | Memory/Learned | 19.82 | [0.87, 0.96] | 6.27 |
|  | Memory/Famous | 3.64 | [0.58, 0.82] | 1.15 |
| CB | Perception/Unfamiliar | 5.96 | [0.56, 0.63] | 1.89 |
|  | Memory/Learned | 14.13 | [0.82, 0.94] | 4.47 |
|  | Memory/Famous | 4.06 | [0.57, 0.73] | 1.28 |
| SA | Perception/Unfamiliar | 12.66 | [0.63, 0.69] | 4.00 |
|  | Memory/Learned | 13.66 | [0.73, 0.83] | 4.32 |
|  | Memory/Famous | 4.22 | [0.56, 0.71] | 1.33 |

Note: CI = confidence interval.

### Reconstruction consistency across participants

To assess the consistency of perceptual reconstructions across our three main participants, we correlated image-based accuracies of the 57 unfamiliar faces across pairs of participants in Experiment 1. This analysis found significant Pearson correlations in every case (NC-CB: r(55) = .66; NC-SA: *r*(*55*) = .65; CB-SA: r(55) = .61; all ps < .001). Similar results were found by correlating experimental-based estimates based on group-averaged data of naïve participants in Experiment 2 (NC-CB: r(55) = .54; NC-SA: r(55) = .62; CB-SA: r(55) = .62; all ps < .001).

## Discussion

The current work aims to achieve image reconstruction from both perception and memory on the basis of behavioural data. Notably, this work demonstrates, for the first time to our knowledge, that the appearance of facial identity can be successfully extracted and reconstructed from an individual’s long-term memory. This demonstration evinces a number of theoretical and practical implications, as discussed below.

First, our empirical data were generated by appeal to a simple, intuitive task, requiring participants to judge the similarity between a current stimulus and a face recalled from memory, in order to derive a hybrid perception-memory face space. The twofold success of perception and memory-based reconstructions relying on this construct provides direct evidence for its psychological validity and furthermore, for the close integration of perception and memory in visual processing. Specifically, the ability to use perceptual features extracted from face stimuli to reconstruct the appearance of faces recalled from memory is consistent with the hypothesis of visual representations shared across perception and memory. Although the present work focuses on faces as a visual category, it is likely that this integration extends to other categories such as objects and scenes^26–28^. According to this viewpoint, therefore, it is useful to consider perception and memory as highly interactive and mutually constraining cognitive processes. Converging with this idea, neuroimaging work has established that visual imagery and perception rely on overlapping neural resources^29–31^ and that the degree of overlap may determine the strength of vividness associated with mental imagery^32^. Moreover, there has been an accumulation of data from neuropsychological and neuroimaging studies to suggest that brain regions typically associated with long-term memory processing (i.e., the medial temporal lobe structures) also play a crucial role in perception^33–37^. Thus, as evidence for the integration of different types of visual processing comes into sharper focus, future work will be needed to elucidate the precise neural mechanisms by which visual representations are recruited and (re)shaped by perception, imagery, and memory processes.

Second, regarding the nature of visual representations, the present work provides evidence that perception and memory share pictorial content, as needed for reconstruction purposes, rather than only higher-level semantic content. Notably, our results show that face representations contain sufficient pictorial detail to support image reconstruction even of faces retrieved from long-term memory. The accuracy of these reconstructions was estimated by the same participants who provided the original reconstruction data as well as by an independent group of naïve participants, and in all cases, accuracy estimates were above chance. Interestingly, the level of reconstruction accuracy for newly-learned faces surpassed that corresponding to famous faces and even to viewed unfamiliar faces. This outcome is likely due, as intended, to the extensive familiarisation of our participants with a small set of face images that aimed to facilitate access to relevant facial features in the memory task. Arguably, familiarisation would not only allow a visually richer experience with faces recalled from memory but also could refine the representation of these faces over time by allowing the participants to zero in on features diagnostic for identification and encode such features preferentially^38^. Therefore, the level of relevant visual detail associated with certain memories seems comparable if not superior to that provided by perceptually available stimuli. At the same time though, we note that low-level image properties were controlled in all face stimuli. Further, in the case of famous face reconstruction, representations were not associated with specific images throughout the experiment as participants were required to retrieve the appearance of the famous faces from their own personal knowledge. Hence, our results go beyond low-level visual processing and, likely, speak to intermediate-level visual representations of facial identity^39–41^.

Third, reconstruction was carried out with the aid of facial features synthesised directly from experimental data rather than predefined, pre-selected ones, such as those extracted from face images via principal component analysis or independent component analysis^15,42^. Specifically, we appealed to a technique akin to reverse correlation to derive facial features from perceptual data and then used such features for image separately for perception and memory-based reconstruction. This aspect of the procedure is significant in that it not only serves immediate reconstruction purposes but also, it helps to clarify the featural basis of perceptual/memory representations and to validate their psychological plausibility through their use in image reconstruction.

Fourth, while reconstruction, as a general approach, was initially designed to accommodate neural data^43^ our work exploits, instead, the structure of behavioural data. This fact is significant in that behavioural-based reconstructions may be more robust than their neural-based counterparts^12^ and could also serve to validate the latter. The current work, therefore, not only helps to establish the utility of behavioural-based reconstruction to memory research as an autonomous method but also highlights its value in relating neural and behavioural aspects of memory processes.

Of note, our investigation targeted reconstructions of face images able to support identification using a sample of homogeneous face stimuli (i.e., frontal views of adult Caucasian male faces). We reasoned that if reconstructed images can be discriminated from each other even within a face dataset exhibiting high inter-similarity it would be considerably easier to achieve successful reconstruction in the presence of broader visual cues introduced by race or gender with a more heterogeneous dataset. However, the validity of this assumption and its precise extent remain to be investigated.

Further, our reconstructions tended to capture primarily low and medium spatial frequency information, typical of classification images derived through reverse correlation^22,23^. Specifically, the comparison of a visual template to a stimulus is prone to spatial uncertainty as the observer applies the template over a range of spatial locations in the stimulus, leading to the smearing of the signal over the region of uncertainty and thus, to blurred CIMs^44^. Here, CIM features appeared to encode extensive shape and surface information but much less, if any, high-frequency fine-grained textural information. However, this is not a limitation in the present case: while our face recognition system exhibits considerable flexibility^45^, we tend to rely on a narrow band of low spatial frequencies for face identification^46,47^ optimal for exploiting the statistical properties of facial images^48^. Thus, since the aim of reconstruction is not to produce a photographic replica of a given stimulus but rather to extract and to visualise the representational content supporting recognition, it appears that the methodology deployed here is largely successful in this respect.

It is important to note that in order to maximise our ability to validate the current research paradigm, participation was restricted in Experiment 1 to individuals with high levels of recognition performance (see Screening, Supplementary Information). Thus, further research will be required to confirm the general applicability of our approach to a broader population of healthy adults and, potentially, of individuals with perceptual/memory-based recognition deficits, as well as to clarify the precise nature of the personal experience and individual characteristics that facilitate successful retrieval of visual information for reconstruction purposes.

Crucially, the consistent success of reconstruction across conditions and across participants in the current study speaks favourably regarding the more general applicability of the present approach and its ability to open up new paths for theoretical and applied investigations. For instance, our approach can be easily directed at testing specific hypotheses regarding face space structure^18,49^, the nature of configural processing^50,51^ or the developmental trajectory of face recognition and its representational basis^52,53^. Further, optimised versions of the method above can serve as a basis for forensic applications. For instance, automated ‘sketch artists’ relying on judgments of facial similarity, instead of verbal descriptions, could provide a robust complement to current strategies for depicting and visualising the face of a person of interest.

## Methods

### Experiment 1 – facial image reconstruction

#### Participants

We sought to assess our approach separately with three participants (NC, Caucasian female, 22 years; CB, Caucasian female, 21 years; SA, Asian male, 26 years) selected based on their performance in several screening tests. On passing our eligibility criteria (see Screening, Supplementary Information), these participants completed four 1-hour experimental sessions on separate days over the course of at most two weeks. All participants had normal or corrected-to-normal vision and no history of neurological or visual disorders. Informed consent was obtained from all participants. All procedures were carried out in accordance with University of Toronto Research Ethics Guidelines and were approved by the University of Toronto Research Ethics Board.

#### Stimuli

Sixty unfamiliar face images selected from four databases: FERET^54,55^, FEI^56^, AR^57^, and Radboud^58^, along with thirty images of famous individuals (i.e., media celebrities) from publicly available sources were selected to display front views of Caucasian males with a neutral expression. All images were cropped, spatially normalised, and colour-normalised for mean values and contrast separately in each CIEL*a*b* colour channel.

Next, from our pool of 60 unfamiliar faces, three were selected to serve as targets for experimentally-controlled face familiarisation and learning (see Novel face learning, Supplementary Information). Also, from our pool of 30 famous individuals, three images were selected for each participant based on their familiarity with the individuals depicted by these images (i.e., at least 5 on a 1-7 familiarity scale for each famous individual) – the remaining famous face images were eliminated from further testing. Of note, while all participants were tested with the same triplet of learned faces for reconstruction purposes, different triplets of famous faces were used for each participant depending on their relative familiarity with different celebrities.

#### Experimental procedures

Data intended for reconstruction purposes were collected with the aid of several pairwise similarity-rating tasks. Specifically, participants performed a perception-based task with unfamiliar faces, and two memory-based tasks, for learned faces and for famous faces, respectively (Fig. 1a).

In the perception-based task, each trial started with a centrally-presented fixation cross (500 ms), followed by a pair of face images presented side by side for 2000 ms against a dark background. Each face subtended an angle of 2.6° × 4° from 90 cm and was displaced 2.4° from the center of the screen. Participants were asked to rate the similarity of the two faces on a 7- point scale by pressing a corresponding number key. The left/right location of the images was counterbalanced and each face was paired with every other face exactly once, leading to 1596 trials divided equally over 14 blocks.

In memory-based tasks, participants were first instructed to recall and hold in memory one of three learned faces or, alternatively, one of three famous individuals. In each trial, a 600 ms central fixation cross was replaced by one of the 57 unfamiliar faces for 400 ms. Participants rated the similarity between the presented face and the recalled face on a 7-point scale, and a 100 ms white-noise mask appeared at the center of the screen as soon as a response was recorded. Each learned/familiar face was paired with every other unfamiliar face once (171 trials per memory-based task, spread over 9 blocks). Of note, participants were not exposed to any images of the three famous faces during this testing, nor did they encounter such images outside of the lab via other means (e.g., media), as they confirmed at the end of the experiment. In contrast, the learned faces were presented at the beginning of each memory-based block so as to refresh their memory of these faces.

For all tasks, trial order was randomised and practice trials were provided at the beginning of each session. Data collection relied on Matlab with the aid of Psychtoolbox 3.0.12^59,60^.

#### Reconstruction procedure

Our approach broadly followed that of Nestor et al.^12^ with the main difference that, first, reconstruction was performed separately for each participant rather than at the group level and second, perception-based reconstruction was accompanied by its memory-based counterparts. Briefly, the method involved: (i) computing a confusability matrix that contained the average pairwise similarity of *n-1* unfamiliar faces (Fig. 1b); (ii) estimating a 20-dimension face space by applying metric MDS to the confusability matrix of each participant and normalising each dimension by z-scoring (Fig. 1c); (iii) deriving CIM’s by deploying, separately for each dimension, an analogue of reverse correlation that computes a weighted average of face images proportionally with their coordinates; (iv) assessing CIM significance through a pixelwise permutation test (i.e., by randomising images with respect to their coordinates on each dimension and by recomputing CIM’s for a total of 10^4^ permutations; pixelwise two-tailed t-test; FDR correction across pixels: q<0.1); (v) projecting a target face (image *n* in Fig. 1b, c) in the existing face space based on its similarity with the *n-1* faces, and (vi) reconstructing the appearance of the target face through a linear combination of significant CIM’s added onto an average face image derived from the linear combination of the original *n-1* faces (Fig. 1e).

Importantly, the procedure above enforces non-circularity by excluding the target face from the estimation of the CIM’s that enter its reconstruction. Specifically, memory-based reconstruction used the 57 unfamiliar faces to estimate face space features while the learned/famous faces provided the reconstruction targets. Similarly, perception-based reconstruction utilised a leave-one-out schema by using 56 unfamiliar images at a time to derive facial features while the remaining face was the reconstruction target (see Face space and facial feature derivation, Image reconstruction procedure in Supplementary Information).

#### Image-based evaluation of reconstruction results

Objective image-based reconstruction accuracy was measured as the pixelwise similarity of reconstructed images relative to the target faces. Specifically, accuracy was estimated as the percentage of instances for which a reconstruction image was closer, via an Euclidean metric, to its target than to any other alternative image. For perception-based reconstruction alternative images were provided by all unfamiliar faces other than the target; similarly, for memory-based reconstruction of learned faces the alternatives to any target were provided by the other two learned images. Such estimates were not computed for famous face reconstruction since no corresponding visual stimuli were presented during the main part of the experiment.

Next, reconstruction accuracies were averaged across all faces, separately for each participant, and tested against chance (50%) using a one-sample t-test. Notably, significance testing was conducted solely for perception-based reconstruction though, and not for memory-based reconstructions, due to the small sample size (and, also, due to the absence of relevant estimates, for famous faces). Hence, to provide a more thorough evaluation of reconstruction results and to complement the image-based assessment above a second experiment was conducted as follows.

### Experiment 2 – experimental evaluation of reconstruction results

#### Participants

In addition to our three participants above, 30 other naïve participants (16 female; age: 18-32 years) were recruited for this experiment – we deemed a sample of this size would suffice for the purpose of capturing effects as robust as those found with the image-based procedure described above. Each session took one hour to complete - for NC, CB and SA, this additional session was conducted within two days of completing Experiment 1.

#### Experimental procedures

In three separate conditions (corresponding to perception, memory-learned, and memory-famous face reconstructions), participants systematically evaluated the similarity between a reconstructed image and two potential targets using two-alternative forced-choice testing. For perception-based and for memory-based reconstructions of learned faces, participants were shown a reconstructed image at the top of the screen alongside two images at the bottom (i.e., a target and a randomly selected foil from the remaining 57 unfamiliar faces or the other 2 learned faces). Participants then selected, via a button press, the bottom image that was the most similar to the top one. In contrast, for memory-based reconstructions of famous faces, each reconstructed image was paired with two names (the target plus a randomly selected name for one of the other 2 famous faces) and participants had to judge which of the two named individuals was closest in appearance to the reconstruction (Fig. 1f). Each trial lasted 2s (perception) or 3 s (memory), and a 100 ms white-noise mask appeared at the location of each stimulus following a response. For the perception-based condition, each reconstructed image was presented 8 times (4 blocks of 114 trials) whereas for the two memory-based conditions, each reconstructed image was presented 36 times (2 blocks of 54 trials).

Of note, NC, CB and CA evaluated their own reconstructions whereas each new participant assessed reconstructions derived from a single participant in Experiment 1, depending on their relative familiarity with different famous face triplets (as used with NC, CB or SA).

Experimental-based reconstruction accuracy was next computed as the percentage of instances for which a reconstructed image was matched correctly with its target. Accuracies were averaged across reconstructions, separately for each condition, and then compared against chance (one-sample two-tailed t-test across participants against 50% accuracy). Mean accuracies were also analysed using a mixed-design two-way analysis of variance (3 within-participants reconstruction types x 3 between-participants reconstruction source: NC, CB or CA).

## Acknowledgments

This work was supported by the Natural Sciences and Engineering Research Council of Canada (AL, AN) and a Connaught New Investigator Award (AN). Portions of the research in this paper use the FERET database^54,55^ of facial images collected under the FERET program, sponsored by the DOD Counterdrug Technology Development Program Office.

## Author Contributions

C.-H. Chang, A. C. H. Lee, and A. Nestor developed the study concept, designed the experiments and wrote the manuscript; C.-H. Chang performed data collection; C.- H. Chang and D. Nemrodov performed data analysis.

## Additional Information

### Supplementary Information

Competing Financial Interests: The authors declare no competing financial interests.

